# A simple spatially explicit neutral model explains range size distribution of reef fishes

**DOI:** 10.1101/238600

**Authors:** Adriana Alzate, Thijs Janzen, Dries Bonte, James Rosindell, Rampal S. Etienne

**Author notes:** Joint last authors (James Rosindell, Rampal S. Etienne).

## Abstract

**Aim:** The great variation in range sizes among species has fascinated ecologists for decades. In reef-associated fish species, which live in fragmented habitats and adopt a wide range of dispersal strategies, we may expect species with greater dispersal ability to spread over larger ranges. However, empirical evidence for such a positive relationship between dispersal and range size in reef fishes remains scarce. Here, we unveil the more nuanced role of dispersal on the range size distribution of reef associated fishes using empirical data and a novel spatially explicit model.

**Location:** Tropical Eastern Pacific

**Major taxa studied:** Reef-associated fishes

**Methods:** We estimated range size distributions for six different guilds of all reef-associated fishes with different dispersal abilities. We used a one-dimensional spatially explicit neutral model, which simulates the distribution of species along a linear coastline to explored the effect of dispersal, speciation and sampling on the distribution of range sizes. Our model adopts a more realistic gradual speciation process (protracted speciation) and incorporates important long distance dispersal events with a fat-tail dispersal kernel. We simulated our model using a highly efficient coalescence approach, which guarantees the metacommunity, is sampled at dynamic equilibrium. We fitted the model to the empirical data using an approximate Bayesian computation approach, with a sequential Monte Carlo algorithm.

**Results:** Stochastic birth, death, speciation and dispersal events alone can accurately explain empirical range size distributions for six different guilds of tropical, reef-associated fishes. Variation in range size distributions among guilds are explained purely by differences in dispersal ability with the best dispersers covering larger ranges.

**Main conclusions:** A simple combination of neutral processes with guild-specific dispersal ability provides a general explanation for both within- and across-guild range size variation. Our results support the theoretically expected, but empirically much debated, hypothesis that dispersal promotes range size.

## Introduction

What is driving the large natural variation in the range size of species (Gaston 2003)? Answers to this long-standing question in macroecology were initially provided by investigating the effects of speciation and extinction processes (Anderson 1985, Gaston & Chown 1999). However, as suggested by Gaston and He (2002), these processes are not sufficient to explain range size distributions in nature, as they only affect the creation, division and removal of ranges. Among the other factors that could influence range size, dispersal ability of individuals is the one with the most important: dispersal is needed for the colonization of new habitats, and for persistence in existing habitats that are suboptimal, where demographic rescue can act to avoid local extinction (MacArthur & Wilson 1967, Brown & Kodric-Brown 1977). Dispersal also promotes gene flow, bringing the genetic variability necessary for adaptation, which is important for successful colonization and ultimately range expansion (Holt & Gomulkiewicz 1996). One group of organisms for which dispersal seems especially important is reef fishes because they live in habitats that are highly fragmented; making the ability to disperse key for habitat colonization, establishment, and range expansion. Despite theoretical expectations predicting a positive relationship between dispersal and range size, empirical evidence for this in reef fishes remains scarce (Lester & Ruttenberg 2005, Ruttenberg & Lester 2015, Mora *et al*. 2012, Luiz *et al*. 2013).

There are many possible explanations for the apparent lack of a positive range size-dispersal relationship; these reflect the many processes that potentially drive range size (reviewed in Gaston 2003) including speciation, local extinction, and range size changes during a species’ lifetime (Webb & Gaston 2000). Firstly, range size is likely to vary with species age (Webb & Gaston 2000), i.e. older species might have attained larger ranges than newly formed species.

Secondly, species range dynamics are affected by biological interactions, eco-evolutionary dynamics and by their behavioral and functional traits (Stahl *et al*. 2014). Thirdly, sampling intensity and detection probability vary across space and across species (Dennis *et al*. 1999, Alzate *et al*. 2014), and such sampling biases could also drive variation in range size. Finally, stochastic events, especially during early life, may bring additional noise to the final range size, making it difficult to find general patterns.

The dispersal component of range size-dispersal relationships is also problematic: dispersal is a complex trait, varying at several life stages, e.g. during departure, transfer and settlement phases (Bonte 2012), in ways that are not easily quantifiable. This may influence the outcome of studies examining the role of dispersal. For example, many studies of dispersal on reef fishes have focused primarily on the larval stage (Lester & Ruttenberg 2005, Lester *et al*. 2007, Mora *et al*. 2012), despite evidence that dispersal also occurs in earlier life stages as eggs and in late life stages as adult fishes (Leis 1978, Kaunda-Arara & Rose 2004, Appeldoorn *et al*. 1994, Addis *et al*. 2013).

Given the complexity of the problem, a promising approach for understanding the drivers of range sizes (in contrast to the many correlative studies) is to model the process, including one or several possible factors affecting range sizes. Although some previous studies have attempted to explain range sizes using colonization-extinction models (Hanski 1982) or population models (Gaston & He 2002), they were not developed to explain variation in range size across many species exploring several factors. Here, we apply a variation of the unified neutral theory of biodiversity and biogeography (Hubbell 2001), originally used to explain other macroecological patterns such as species abundance distributions, species area relationships and beta-diversity. We extend the neutral model to include spatially explicit dynamics and a more realistic speciation process (Rosindell *et al*. 2008, Rosindell *et al*. 2011), both of which we expect to be important for a study of interspecific variation in range sizes. This mechanistic model provides a way to quantitatively assess how dispersal can influence species range size distributions, while at the same time considering other interacting factors, including both sampling and speciation, that are known to affect range size (Gaston 2003). We tested the ability of our model to explain variation in range sizes by comparing its predictions against empirical range size distributions of a complete reef fish assemblage in a well-defined region: The Tropical Eastern Pacific (TEP). We made predictions of range size distributions for each of six distinct guilds with different dispersal characteristics in the early (egg and larval) as well as the later adult life stages. Our model is neutral and so excludes any within-guild niche-based processes and individual differences. Crucially, by applying independent neutral models to each of the six guilds we were able to focus on studying the effects of different dispersal abilities for each guild in isolation from other complicating factors such as environmental preference. With our spatially explicit neutral model, we tested firstly whether range size distributions within guilds of reef fishes can be explained by neutral factors alone and secondly whether variation in range size distribution across guilds can be explained by differences in dispersal ability.

## Methods

### Reef-associated fish data

From the online database “Shorefishes of the Tropical Eastern Pacific - SFTEP” (Robertson & Allen 2016), we collated spatial coordinates of species occurrences (45.860 records) for all bony fishes (575 species) associated to reef habitats reported in the TEP. We used only records inside the TEP region: 24° N (outer coast of California gulf, including all the inner coast) and 4° S (SFTEP, Robertson & Allen 2016).

Reef fish species were classified in six different dispersal guilds according to traits related to dispersal: spawning mode and adult mobility. We classified spawning mode in two types: pelagic and non-pelagic. The differences in this early life history might confer diverse capacities for dispersal (Riginos *et al*. 2011, Leis *et al*. 2013). Pelagic spawners release their eggs in the water column, which are passively transported by water currents until the larvae hatch and are able to better control active swimming (Stobutzki 1997, Leis *et al*. 2013). This increase in the pre-hatching dispersal period might have strong and broader effect on dispersal in the pelagic environment (Leis *et al*. 2013). Contrary to pelagic spawners, for which both the egg and larval phases are pelagic, non-pelagic spawners either attach their eggs to the substrate, are livebearers, or keep their eggs in the mouth or pouch until they hatch. Their larvae usually emerge at larger sizes and are more mature than the larvae of non-pelagic spawners (Wootton 1992, Leis *et al*. 2013), resulting in an early control of active swimming, therefore limiting dispersal (Munday & Jones 1998, Leis 2006, Leis *et al*. 2013). We classified adult mobility following Floeter and colleagues (2004) as low, medium and high. Low adult mobility denotes site-attached species with a restricted home range (< 10m^2^). Medium adult mobility denotes species that are weakly mobile, relatively sedentary, with close association to the substrate and that can be distributed over the entire reef area (< ~1000m^2^). High adult mobility denotes species that are highly mobile with wide horizontal displacement and that occur in the water column (Floeter *et al*. 2004). Mobility for each species was assigned depending on the taxonomical level at which information was reported: species, genus or family adult mobility. In some cases, mobility information was not available, but could be assigned according to the biology of the species, e.g. pearlfishes (Family Carapidae) that are known to live inside the anal pore of sea cucumbers were all classified as having low adult mobility. Information on adult mobility was obtained from several sources (data base in Suppl. Mat). Information on spawning mode was obtained from the SFTEP online database (Robertson & Allen 2016). Pelagic larvae duration, although often used when studying range size of reef fishes, is not known for the majority (69%) of species in the TEP region, making it unsuitable for this study.

### Measuring range size

The range size of each species was calculated using a novel metric, developed for maximizing comparability between simulated and observed range sizes: coastline distance. In contrast with other traditional metrics, e.g. maximum linear distance, latitudinal and longitudinal extent (Gaston 1994), coastline distance does not underestimate or overestimate range size due to the particular spatial configuration of the TEP (Fig. S1). We defined coastline distance as the contour distance (measured using units of 100 km) between the most distant points along the coast line where the species was reported. However, the east and west coast of the Californian gulf are treated as a single coast because the distance between opposing coasts is likely too small to substantially restrict dispersal at similar latitudes (Fig. S1). All distance measurements were calculated in kilometers using the function geodist from the R package gmt (Magnusson 2015) and transformed in relative values, where 100% is the coastline distance between the latitudes 24N and 4S.

### Spatially explicit neural model

We used a one-dimensional spatially explicit neutral model to simulate the spatial distribution of species along a linear coastline. This configuration best reflects the particular geographical distribution of reefs (coral and rocky) in the TEP region: a long coastline with a narrow continental platform. As in the original neutral model (Hubbell 2001), the habitat is saturated (zero-sum dynamics) and the species identity of an individual has no bearing on its chances of dispersal, mortality, reproduction, the initiation of speciation or the completion of speciation (see below). At every time step one individual, chosen at random according to a uniform distribution, dies and is replaced by the newborn offspring of an existing individual determined by a Pareto dispersal kernel:

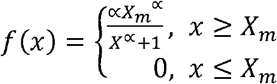

where *X_m_* is a scale parameter (mode) and α is a shape parameter, that changes the distribution from an exponential-like distribution (large value of α) to a very fat-tailed distribution (lower values of α), i.e. many short distance dispersal events are combined with an occasional very long-distance dispersal event. Random samples from the distribution can be calculated using the inverse random sampling formula for the range size *T*:

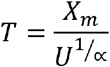

where *U* is a random variate drawn from a uniform distribution between 0 and 1. To separate the effects of the shape of the distribution and the mean dispersal distance (*X*_mean_), we rescaled the Pareto distribution such that *X_m_* = *X*_mean_, i.e. that *X_m_* reflects the mean dispersal distance, and α still reflects the shape (see Suppl. Mat. for full derivation). The Pareto distribution considers the possibility of long distance dispersal, in line with empirical dispersal distributions of reef fishes (Jones 2015).

In contrast to the classical neutral model, we assumed that speciation is a gradual process rather than an instantaneous event (Rosindell *et al*. 2010). When a birth event takes place, an incipient species can form with probability μ; the newborn is still observed in the model as being conspecific to its parent, but if sufficient time passes and descendants of the newborn individual survive, those descendants will be considered a new good species rather than an incipient one. This protracted speciation model entails one extra parameter τ: ‘protractedness’, the number of generations required for an incipient species to become a real species, where one generation means half of the turnover of the community because generations overlap. Both speciation probability and protractedness influence the generation of new species, the true speciation rate is a function of both parameters (μ/1 + τ) as described by Rosindell *et al*. (2010). We simulated the spatially explicit neutral model using a coalescence approach (Rosindell *et al*. 2008), which improves simulation efficiency while guaranteeing the metacommunity is sampled at dynamic equilibrium and thus eliminating the problem of determining an appropriate ‘burn-in time’ for the simulations.

### Model behavior

We explored the effect of dispersal on the distribution of range sizes by running simulations using various dispersal kernels, which differ in their *X*_mean_ and *α* parameter values. We used a linear lattice composed of 100,000 ‘units’ which could be thought of as individual organisms or larger cohorts of individuals behaving in a similar manner (Harfoot *et al*. 2014). We found that larger lattices produce similar results (Fig. S2), but are computationally intractable for parameter fitting exercises that require many successive simulation’ runs. As in the real world not all individuals are sampled, the proportion of sampled individuals (sampling percentage) could therefore affect the observed distribution of ranges. Sampling was performed by randomly choosing individuals along the linear lattice, and only sampled individuals were used to quantify range sizes. Although sample areas along the TEP are not random, sampling in a realistic manner produces virtually similar results as with random sampling (Fig. S3). We examined the effect of dispersal (*X*_mean_ and α), speciation, protractedness and sampling percentage on the distribution of species’ range sizes. As species age is also suggested to be positively related with range size (Gaston 2003), we also explored the effect of interspecific variation in speciation rates on the distribution of range sizes. When speciation rate is high, species are in average younger, thus affecting the final range size distribution.

In our default scenario, we used the following parameter values: *X*_mean_ = 0.02, *α* = 3.0, sampling percentage 5 = 100%, speciation probability μ = 0.0005, protractedness τ = 10. We then performed 5 sets of alternative scenarios, in which either values of *X*_mean_, α, sampling percentage, speciation probability or protractedness were altered. We explored 5 different *X*_mean_ and α values (*X*_mean_ = [2%, 5%, 10%, 20%, 40%], α = [1.5, 2.0, 2.5, 3.0, 3.5]), 5 different sampling percentages (5 = [1%, 5%, 20%, 50%, 100% of all individuals]) and 4 different speciation probability and protractedness values (μ= [5 × 10 ^−2^, 5 × 10 ^−3^, 5 × 10 ^−4^, 5 × 10 ^−5^], *τ* = [0, 10, 100, 1000]).

At the end of our simulations we estimated the range size for each species as the linear distance (which is equivalent to coastline distance in a one-dimensional model) between the most distant points where the species is recorded. The range size was measured in relative terms, relative to the total lattice size. We replicated the simulations 100 times and calculated mean and 95% CI values. Range sizes were transformed to percentages (100 % total size of the linear lattice).

### Model fitting

In order to estimate dispersal (*X*_mean_ and α), sampling, speciation and protractedness values that produced range size distributions matching those of empirical data, we used an approximate Bayesian computation approach, with a sequential Monte Carlo algorithm (ABC-SMC) as described by Toni *et al*. (2009). To assess the similarity between the data and simulation outcomes, we calculated the sum of squares between the inverse cumulative distribution for the simulated and empirical data, based on the differences in both the range size distributions and species richness levels. Progression of the acceptance threshold was modeled as an exponentially decreasing function, where the threshold at iteration *t* of the ABC-SMC algorithm was: 500 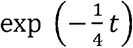. We assumed the following prior distributions for each parameter (on a log_10_ scale, e.g. *U*_10_(0,1) = 10^*U*(0,1)^, where *U* is a uniform distribution), *X*_mean_: *U*_10_(−4, −0.25), α: *U*_10_(0,1), speciation initiation rate: *U*_10_(−5, 0), protractedness: *U*_10_(0,5) and sampling: *U*_10_(−4, 0). Per ABC-SMC iteration, we used 10,000 particles. The ABC-SMC algorithm ran for 20 iterations, or until the acceptance rate dropped below 1 in 1,000,000 proposed parameter combinations. Perturbation of the parameters was performed on a log10 scale, to avoid parameters reaching a negative value. Parameters were perturbed by first taking the log10, then adding a random number drawn from a normal distribution with mean zero and standard deviation 0.05, after which we exponentiated the parameter again. After exponentiation, the parameter values were checked whether they still lay within the prior ranges; if not, the particle was rejected. For each dataset we performed 10 replicate fits.

To assess the accuracy of our inference method, we generated artificial datasets using known parameters, and performed the same ABC-SMC inference procedure as used on the empirical data. If our method is accurate, inferred parameter values should be identical to the known parameters used to generate the artificial data. Artificial data was generated using values for *X*_mean_ of 0.001, 0.01, 0.1 or 0.2, α of 2, 4, 6 or 8, 5 of 0.025 or 0.25, and two different speciation regimes: one with high speciation (0.01) and high protractedness (2500), and one with low speciation (0.001) and low protractedness (25). For each parameter combination we generated 10 artificial datasets. In total we performed (10 × 4 × 4 × 2 × 2) = 640 ABC-SMC inferences to assess accuracy.

The one-dimensionality of our neutral model means the coastline distance metric treats the coast of the TEP as also being one-dimensional (distance is only measured along the coast, not as a birds-flight distance); this maximises the comparability of empirically observed range sizes with those simulated by our one-dimensional, spatially explicit neutral model. In addition, we excluded observations from oceanic islands when quantifying range sizes, again to maximize comparability with simulated ranges. Our model was written in C++ and all post simulation analyses were performed with R, version 3.3.1 (R core team 2016).

## Results (883)

### Range size distribution of reef associated fishes in the TEP

Irrespective of their adult mobility, all three guilds of pelagic spawners have a relatively high proportion of species with large ranges (Fig. 1a). The range size distributions of pelagic spawners are qualitatively similar, with more than 70% of the species having ranges larger than 50% of the maximum possible range or our sampling region. In contrast, the range size distribution of non-pelagic spawners depends strongly on the capacity of adult fishes to disperse. Within the non-pelagic spawners, the lowest dispersive guild has the highest proportion of species with small ranges and the lowest proportion of species with large ranges (Fig. 1a). While more than half of the species with medium or high adult mobility have ranges larger than 80% of the maximum range, for species with low mobility only a fifth of species have ranges larger than 80% of the maximum.

**Fig. 1.**
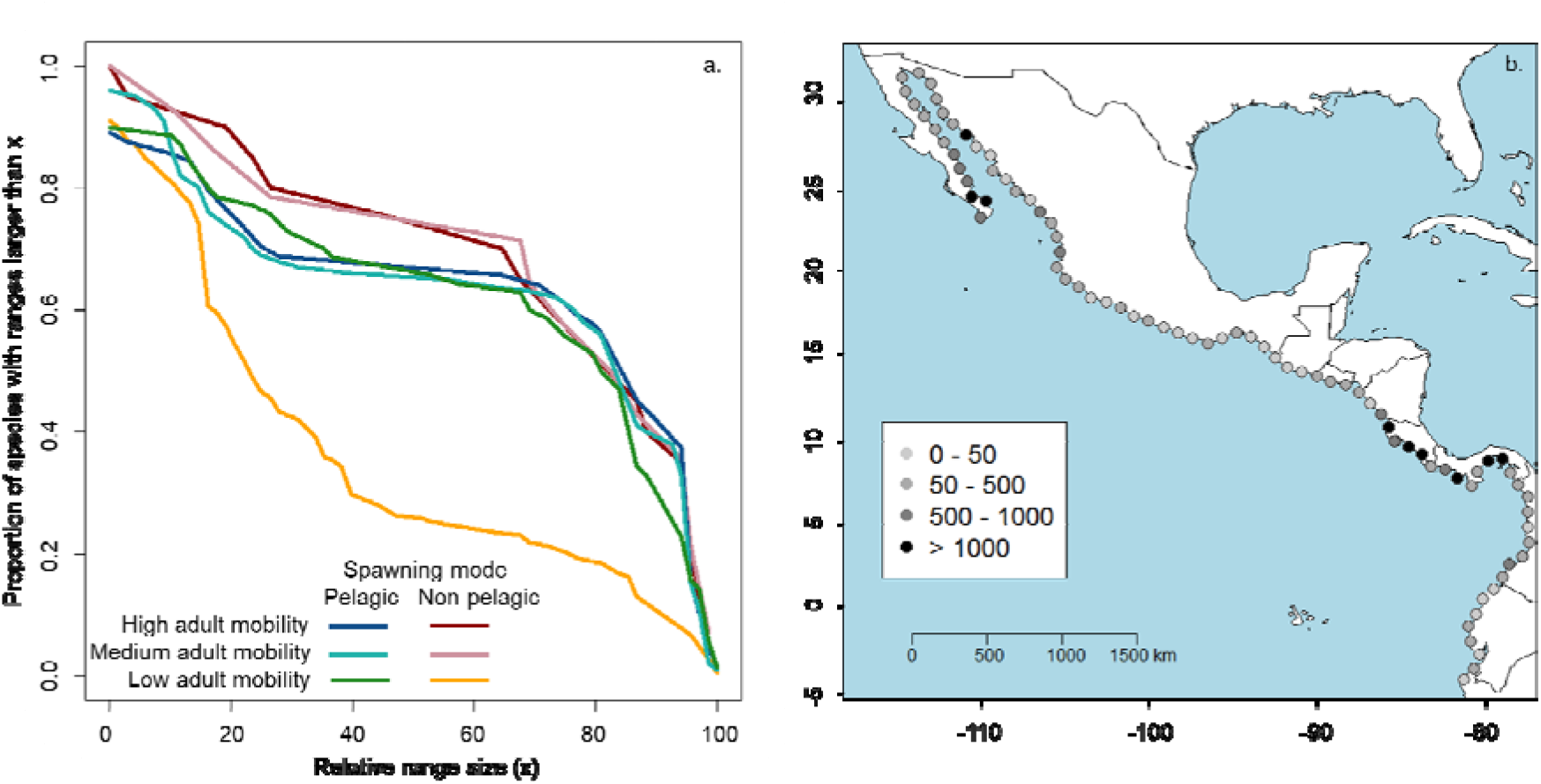
(a) Distribution of range sizes for different dispersal guilds of reef fishes in the Tropical Eastern Pacific (TEP). A guild is defined as a group of species that share the same spawning mode (pelagic and non-pelagic spawners) and adult mobility (low, medium and high). Range size is shown in relative terms, where a range of 100% is the largest range recorded for a species in the TEP. We used coastline distance as the range size metric, which is the distance between the most distant points along the coastline. Individuals from oceanic islands are excluded to be consistent to the one-dimensional nature of the model. Both sides of the California Gulf coastline were shrunk into a single one. The distribution of ranges is shown as cumulative distribution curves, which show the proportion of species (y axis) that attain ranges larger than a given size (x axis). (b) Map showing the sampling intensity along the coastline in the TEP: the number of occurrences recorded at each coastline point spaced by 100 km.

### Spatially explicit neutral model

The strongest effects on the distribution of range sizes are caused by variation in mean dispersal distance (*X*_mean_), speciation rate, and protractedness (Fig. 2). Dispersal (*X*_mean_ and *α*) has a strong effect on the shape of the range size distribution. The contributions of *X*_mean_ and *α* to the effect of dispersal on the range size distribution are not equal however, with the majority of the dispersal effect resulting from *X*_mean_ (Fig. 2a). As *X*_mean_ increases, the proportion of species with large ranges increases as well. In contrast, the shape parameter of the dispersal kernel (*α*) has limited influence over the distribution of range sizes (Fig. 2b). Speciation exerts a strong effect on the distribution of ranges, with a higher proportion of species having a large range size when speciation rate is low. A high speciation rate produces more new species, which initially have small ranges, thus a decrease in the number of species with large ranges, and a (potentially unrealistically) high number of species in total (Fig. 2d). The effect of protractedness is similar to that of speciation, as it modifies the number of species and the rate at which these are created. The higher the protractedness, the longer the time before an incipient species becomes a good species, and as a result fewer species have small ranges (Fig. 2e). Sampling affects the distribution of ranges in a different way to dispersal, speciation or protractedness: a lower sampling effort leads to more species with few individuals and thus a higher proportion of species with apparently small ranges (Fig. 2c).

**Fig. 2.**
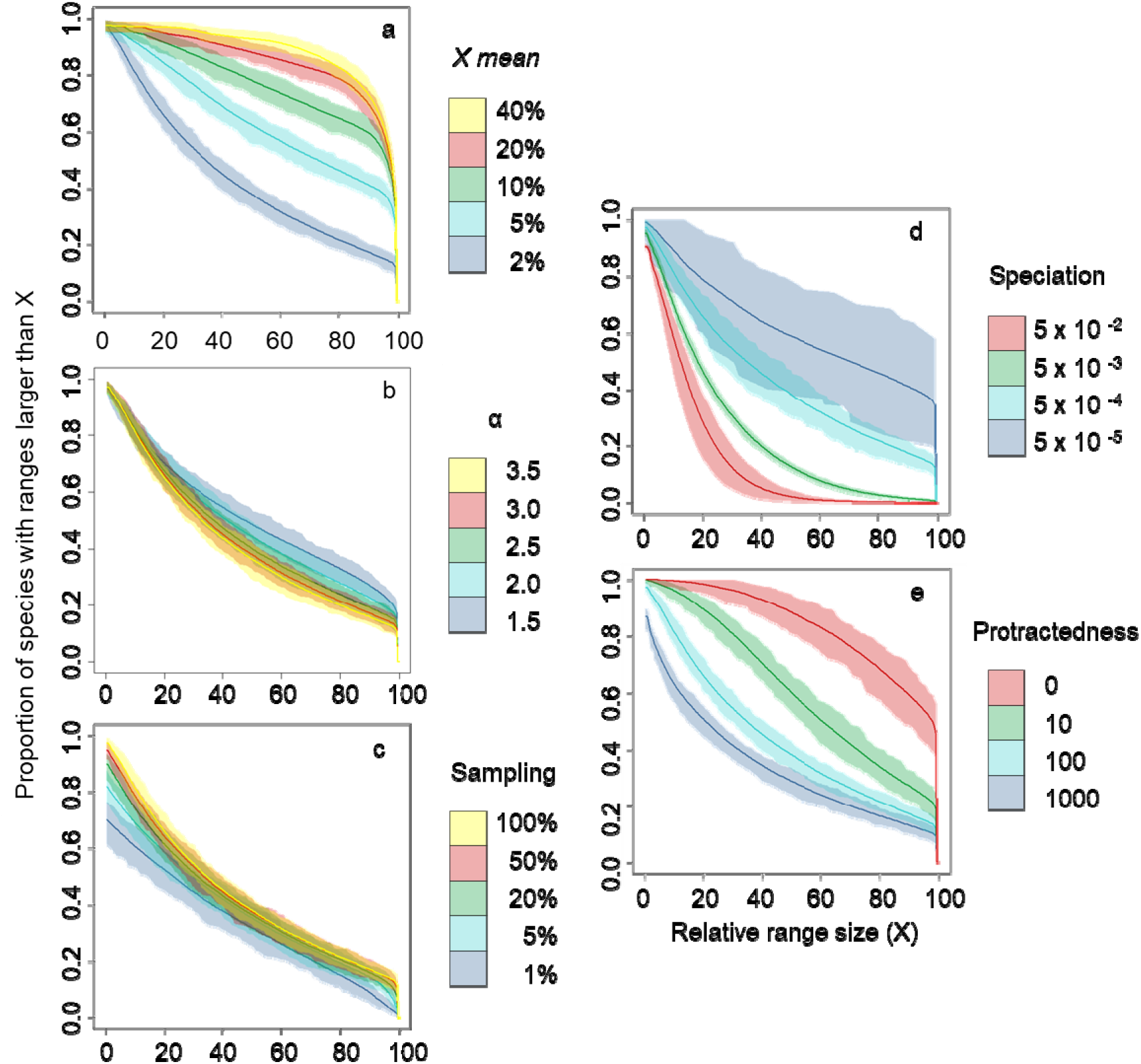
Effect of (a) the mean dispersal distance *X*_mean_, (b) the shape parameter of the dispersal kernel *α*, (c) the sampling proportion, (d) speciation μ, and (e) the time to speciation τ (protractedness) on the distribution of range sizes. Lines show the average value of 100 replicates and the shadows represent the 95% CI. For all simulations, the lattice size was 100,000 individuals. We use one fixed parameter setting, for which only the variable of interest varied: *s* = 100%, *α* = 3.0, *X*_mean_ = 0.02, μ = 0.0005, τ = 10.

Prior to fitting the model to empirical data, we used the ABC-SMC fitting procedure on simulated range size distributions with a known set of parameters (known values for *X*_mean_, α, speciation, sampling and protractedness). We found that posterior distributions of parameter values were generally closely matching the real values (Fig. S4), indicating that our fitting procedure was appropriate for estimating the parameter values of our neutral model. Only in the case of the α parameter (measuring the shape of the dispersal kernel), were estimates were not accurate, likely due the low strength of α in explaining range size variation (see above).

The same fitting procedure on empirical range size distributions, for the six dispersal guilds of reef fishes, showed adequate fit between observed and predicted range size distributions (Fig. 3). Furthermore, in line with expectations, the estimated mean dispersal distances for each guild were largest for the guilds with the highest proportion of large ranges which were pelagic spawners and guilds with high adult mobility as expected. α values were similar for all dispersal guilds (between 3.4 and 4.7). Estimated sampling completeness was lowest for the guilds of non-pelagic spawners with high and medium mobility (0.76 and 0.48% respectively), similarly low for the guild of pelagic spawners (3 – 9%) and very high for the guild of non-pelagic spawners with low adult mobility (38%). Protractedness (the time it takes for an incipient species to become a true species) values were the lowest for non-pelagic fishes, low mobility species (13 generations), while values were intermediate for pelagic spawners (160–730 generations) and highest for non-pelagic spawners with high and intermediate mobility (3500 and 7000 generations respectively). The speciation probability parameter (giving probability for an individual to become a new incipient species) was similar to protractedness being low for pelagic spawners (0.02–0.03), similarly high for non-pelagic spawners with high and medium adult mobility (0.08, 0.06) and the lowest for non-pelagic spawners with low adult mobility (0.0007). See Table S1 for a complete description of the model estimates.

**Fig. 3.**
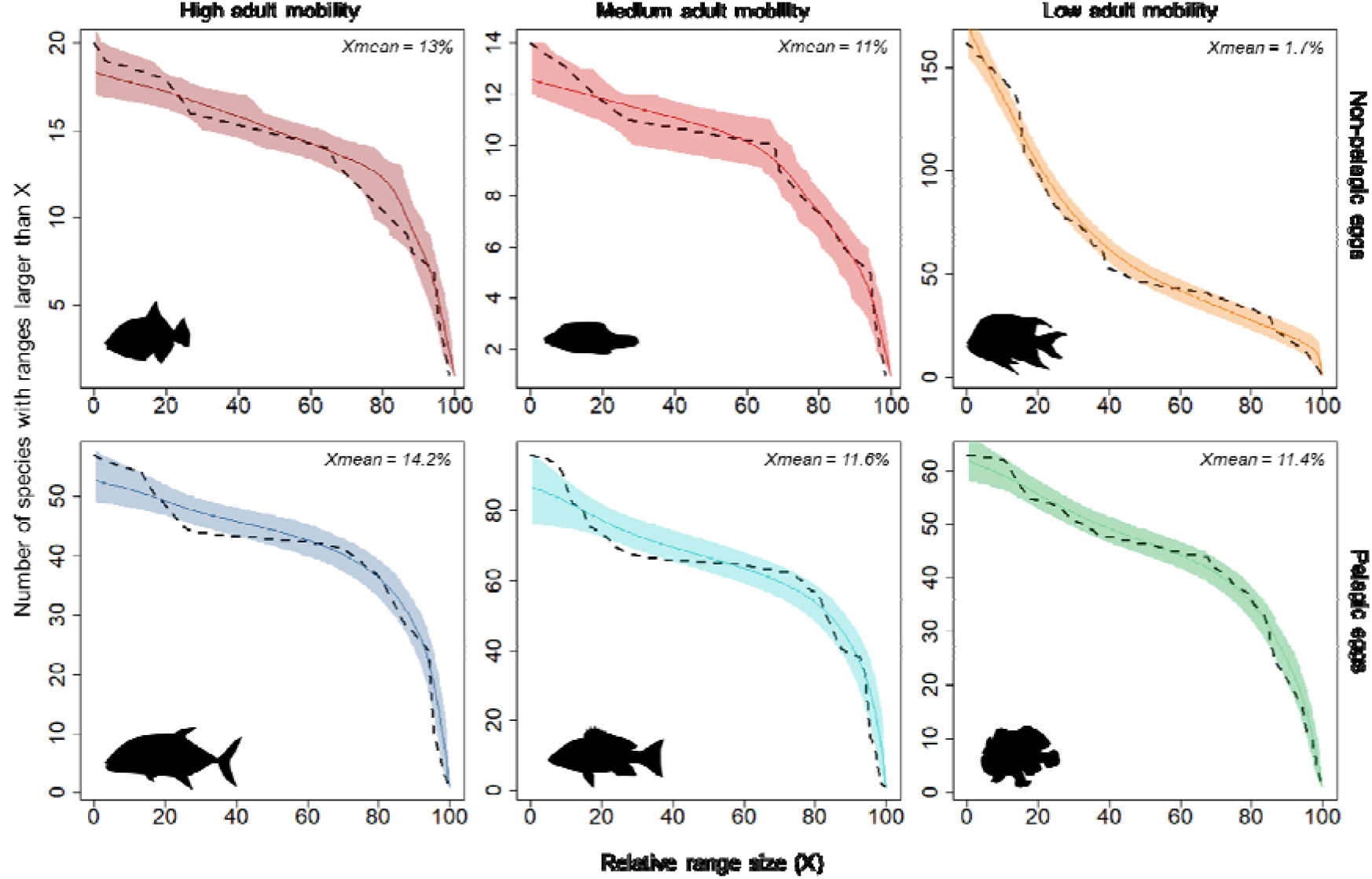
Range size distributions of the best model fitted to each dispersal guild, shown as an inverse cumulative distribution curve. Mean of 5 replicates and 95% CI are shown. Dashed lines represent the empirical data and coloured bands represent the distribution of values in the best fitting model for that guild. Estimated *X*_mean_(mean of >90.000 estimates) are shown per each dispersal guild.

For two dispersal guilds (pelagic spawners with high and medium adult mobility), our neutral model could not fully explain the bimodality in their range size distribution. This mismatch was strongest for pelagic spawners with medium adult mobility (Fig. 3). To explore what caused these mismatches, we performed further analyses, in which we plotted the distribution of ranges for fishes that are endemic to the TEP and one for the non-endemics (following Robertson & Allen 2016). The distribution of ranges in the TEP for these two groups showed differences for all guilds, but especially for the guild of pelagic spawners with medium mobility (Fig. 4). In this case, the bimodality does not appear in either endemics or non-endemics when separated, the combination of these two different distributions thus explains the observed bimodality in the overall distribution.

**Fig. 4.**
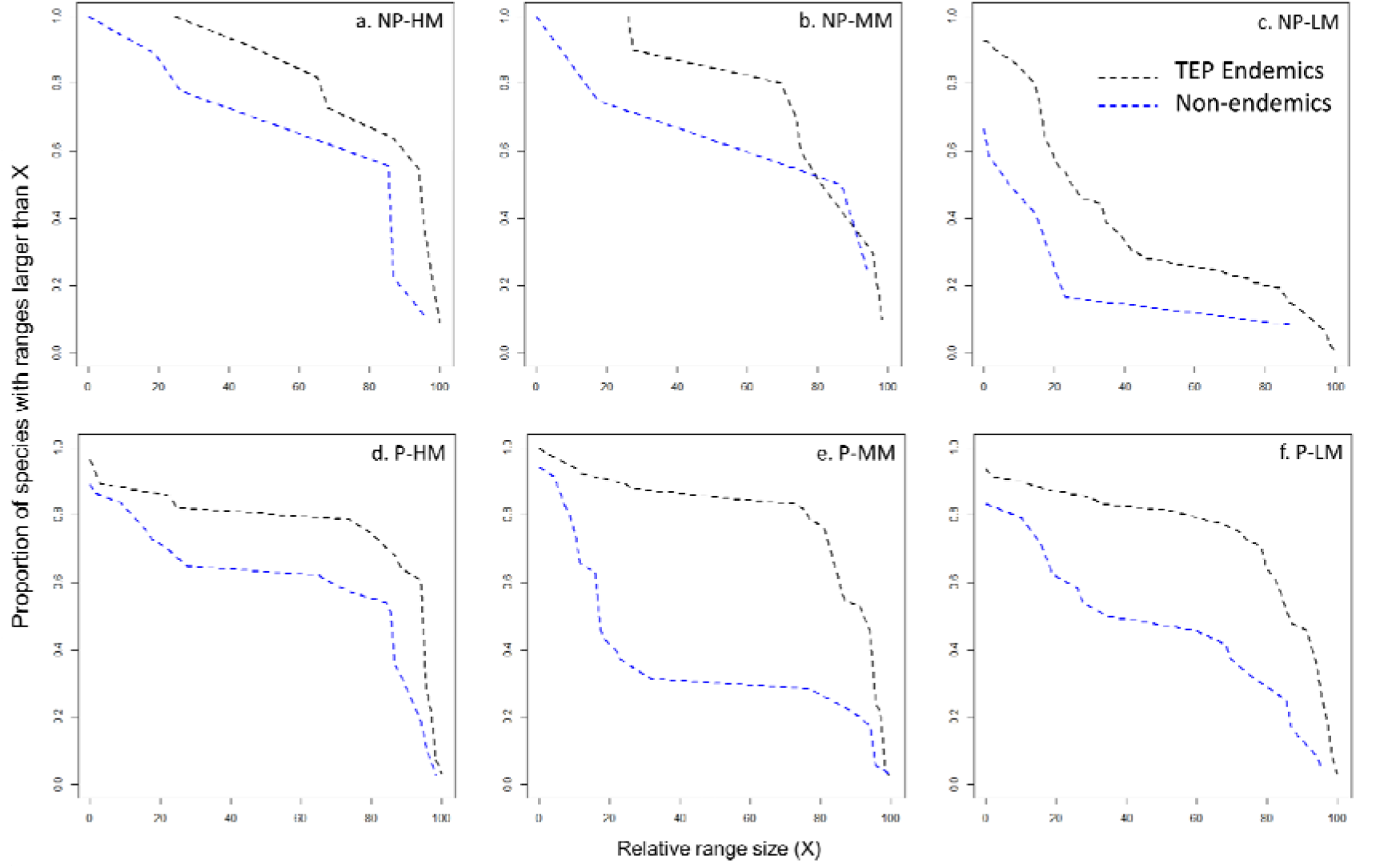
Empirical range size distributions (as inverse cumulative distribution curves) for each dispersal guild. The data shown separately for species that are endemics to the TEP and for TEP non-endemic species.

## Discussion

For decades, macroecologists have tried to understand the large variation in range sizes across species. Using a new approach comprised of several spatially explicit neutral models, we have shown that range size variation can emerge from stochastic birth, death, speciation and variable dispersal abilities. Due to the mixed results of other studies, the importance of dispersal ability in explaining range size variation has often been questioned (Lester & Ruttenberg 2005, Luiz *et al*. 2013, Ruttenberg & Lester 2015). Here, we show that dispersal is really an important factor shaping the range size distribution of species, but that our detailed analyses were required to see this. For example, a study of only species with pelagic eggs may not have revealed any clear effect of dispersal. High dispersal produces distributions with a large proportion of species with large ranges, whereas low dispersal produces a large proportion of small ranged species, consistent with a positive relationship between dispersal and range size. Our model, however, also shows that range size variation can be large within dispersal guilds, as dispersal only affects the *probability* to have large or small ranges. Thus, although low dispersal produces distributions with a large proportion of small ranges, there are also some species with large ranges, and vice versa for high dispersal. This also helps explain why it has been challenging for empirical studies to show clear links between dispersal ability and range size: for each level of dispersal ability, a large variation in range sizes is still possible. Our neutral model predicts range size distributions with a close fit to the empirical distributions for six different dispersal guilds of reef fishes in the TEP, and for each guild estimated mean dispersal distance was in line with expectations, indicating that despite their simplicity, neutral models still capture the most important processes for driving range size variation within such guilds. Importantly, the neutral models we used were originally developed to understand other macroecological patterns (Hubbell 2001), and thus can be seen as an independent mechanistic tool, rather than a phenomenological construct tailored to fit one pattern only.

Although our models generally fitted empirical range size distributions adequately, there were some exceptions. Within guilds of pelagic spawners with high and medium adult mobility, range size distribution tended to be bimodal, something that could not be explained by neutral processes alone. We found that this bimodality primarily resulted from the combination of two different background distributions: TEP endemics vs. TEP non-endemics, with the endemics generally having larger ranges within the TEP. We hypothesize that the former have had a longer time to increase their ranges in the region whilst the latter are biased by including the edges of many wider ranged species that mostly occupy areas outside the TEP. We also found that the range size distribution of non-endemic, pelagic spawners with medium mobility was bimodal (Fig. 4e). A possible explanation is that this is due to their origin, with some species coming originally from temperate regions (North and South America), and others from tropical areas outside the TEP. We conjecture that the majority of species with large ranges are trans-Pacific species, already adapted to tropical conditions. In contrast, 22 out of the 24 species with very small ranges come from temperate regions, and it is likely that their adaptations to a temperate climate and asymmetrical dispersal made these species less able to expand their ranges into areas with more tropical conditions (Holt 2003). In fact, species coming from the temperate north do not go down to the south and vice versa, whereas transpacific species are well distributed along the coast (Fig. S5).

Our results showed that in addition to dispersal, speciation and sampling intensity can also play an important role in shaping the distribution of range sizes. When sampling effort was low, only a single individual was detected for many species (hence they were treated as singletons, even if more individuals were present but not observed), leading to a high proportion of species with very small ranges. The proportion of species with small ranges also increased when speciation rates were high, or when speciation was a fast, non-gradual process (low protractedness). In these cases, new species emerged continuously with low abundance and restricted range. This outcome is in line with hypotheses attempting to explain why range sizes in the tropics are usually smaller than in temperate regions such as ‘Rapoport’s rule’ (Rapoport 1982), which proposes that higher speciation rates in the tropics have caused this pattern (Stevens 1989). Future empirical studies may potentially provide better tests of the validity of our model outcomes. For instance, our predictions of how observed range size distributions change when communities are increasingly intensively sampled, leading to larger ranges as second conspecific individuals are seen for many singleton species.

While we could explain range size distributions using neutral models within guilds, average range size varied across guilds, and observed species characteristics: both differences in adult mobility and spawning mode. The estimated dispersal abilities from our models suggest that differences in average range size are strongly influenced by dispersal. Consistent with previous studies on neutral models with guild structure (using predictions for abundance instead of range size, Janzen *et al*. 2015, Aduse-Poku *et al*. 2017), our results show that while community dynamics within guilds may be captured by a neutral model, across guilds niche-based processes drive variation in range size. Neutral theory was originally proposed to describe community assembly within guilds (Hubbell 2001), and our results are consistent with this philosophy. We take the concept further however, and show that across guilds, niche-based processes, in this case differing dispersal strategies, play a larger role in driving ecological patterns.

We have shown here how variation in range size across species can be explained by a combination of neutral processes and guild-specific differences in dispersal. Our findings thus make substantial progress towards settling a long-standing debate about the underlying causes of variation in range size, and the role of dispersal in this pattern.

## Acknowledgments

AA was funded by the Ubbo Emmius Fund and by BelSpo IAP project ‘SPatial and environmental determinants of Eco-Evolutionary DYnamics: anthropogenic environments as a model’. TJ is grateful for use of the computational cluster ADA of the Max Planck Institute for Evolutionary Biology, Plön, and the use of the computational cluster Peregrine, of the University of Groningen. JR was funded by fellowships from the Natural Environment Research Council (NERC) (NE/I021179, NE/L011611/1). Through JR, this study is a contribution to Imperial College’s Grand Challenges in Eco-systems and the Environment initiative. DB and RE received support from the FWO research community EVENET and FWO project G.018017.N. RSE received support by a VICI grant (865.13.003) from the Netherlands Organisation for Scientific Research (NWO).

